# Tumoroscope: a probabilistic model for mapping cancer clones in tumor tissues

**DOI:** 10.1101/2022.09.22.508914

**Authors:** Shadi Darvish Shafighi, Agnieszka Geras, Barbara Jurzysta, Alireza Sahaf Naeini, Igor Filipiuk, Łukasz Rączkowski, Hosein Toosi, Łukasz Koperski, Kim Thrane, Camilla Engblom, Jeff Mold, Xinsong Chen, Johan Hartman, Dominika Nowis, Alessandra Carbone, Jens Lagergren, Ewa Szczurek

## Abstract

Spatial and genomic heterogeneity of tumors is the key for cancer progression, treatment, and survival. However, a technology for direct mapping the clones in the tumor tissue based on point mutations is lacking. Here, we propose Tumoroscope, the first probabilistic model that accurately infers cancer clones and their high-resolution localization by integrating pathological images, whole exome sequencing, and spatial transcriptomics data. In contrast to previous methods, Tumoroscope explicitly addresses the problem of deconvoluting the proportions of clones in spatial transcriptomics spots. Applied to a reference prostate cancer dataset and a newly generated breast cancer dataset, Tumoroscope reveals spatial patterns of clone colocalization and mutual exclusion in sub-areas of the tumor tissue. We further infer clone-specific gene expression levels and the most highly expressed genes for each clone. In summary, Tumoroscope enables an integrated study of the spatial, genomic, and phenotypic organization of tumors.

## Main

Tumor evolution proceeds by the accumulation of mutations, resulting in the emergence of distinct cancer cell subpopulations, called *clones*, characterized by their genotypes. The spatial distribution of these clones may vary drastically across tumor tissue. This genetic and spatial tumor heterogeneity are the two key determinants of patient prognosis, survival, and treatment [1–3]. Characterization of the phenotypic heterogeneity of tumors, i.e., linking the potential differences between expression profiles of clones and their spatial distribution has up to now remained an uncharted territory.

The vast majority of studies investigate intra-tumor heterogeneity based on bulk DNA sequencing (DNA-seq) or single-cell DNA-seq (scDNA-seq) data [4, 5]. Unfortunately, bulk DNA-seq measures a mixture of millions of cells from different tumors and healthy cells and thus provides only aggregated information of variant allele frequencies. There are several approaches for clonal deconvolution of bulk DNA-seq data, reconstructing the clone genotypes, the frequencies of the clones, and their phylogenetic relationships [6–11]. More recently, several methods for identifying clonal evolution from mutations found in scDNA-seq [12] or from combined bulk and scDNA-seq [13–15] were proposed. Despite recent technological advances [16], scDNA-seq remains much more laborious, more inaccurate, and less affordable than the highly established bulk DNA-seq [17]. Unfortunately, both bulk and scDNA-seq require tissue disaggregation and thus lose spatial information. As methods based on DNA-seq, they cannot be used to elucidate the phenotypic heterogeneity.

Localization of cancer clones has up to now only been obtainable using multi-region single-cell or bulk DNA-seq, combined with computational clone inference for each region [18–21]. This approach, however, is coarse-grained, as each of the regions is itself a bulk sample and is composed of multiple clones with an unknown position in the tissue. Unfortunately, currently there exists no experimental approach for large-scale sequencing of the DNA of single cells *in situ*. However, the recent technology of high-resolution spatial transcriptomics (ST) offers spatially-resolved RNA sequencing (RNA-seq) of mini-bulks of only 1-100 cells, localized in spots of an ST array [22, 23]. Thereby ST enables an analysis of spatial gene expression patterns across the analyzed tissue. Although the resolution of ST is orders of magnitude higher than multi-region bulk sequencing, it still provides only an aggregated signal for mixtures of cells. Moreover, since ST is an RNA-seq protocol and does not have single-cell resolution, it is non-trivial to infer the genotypes of clones at the spots. Finally, single tumor cell phenotypes are widely studied using scRNA-seq, but since the DNA of these cells is not usually measured, the phenotypes are not assigned to the cancer clones. In summary, there exists no state-of-the-art approach for the study of tumor genetic, spatial and phenotypic heterogeneity in a high, close to cellular resolution.

To address this issue, we propose Tumoroscope, a probabilistic graphical model that exploits variant information in ST reads, genotypes of clones reconstructed from bulk DNA-seq and tumor regions and cancer cell counts annotated in hematoxylin and eosin-stained (H&E) images to deconvolute the clonal composition of each localized spot in the tumor sample. On top of that, we devise a regression model for inference of gene expression profiles of the clones. After validating Tumoroscope on simulated data, we set out to answer key questions about co-localization and mutual exclusion patterns of the spatial arrangement of clones and their phenotypes in a newly generated breast, and a previously published prostate cancer dataset [24]. Our approach enables high-resolution spatial mapping and infers gene expression profiles of the clones in the tumor tissue, opening novel avenues in the study of spatial, genetic, and phenotypic heterogeneity of tumors.

## Results

Tumoroscope is a comprehensive probabilistic framework for mapping cancer clones across tumor tissues based on integrated signals from H&E stained images (Fig. 1a), spatially-resolved transcriptomics (Fig. 1b), and bulk DNA-seq (Fig. 1c). The data preprocessing pipeline starts with a two-staged analysis of the H&E-stained image of the tissue (Fig. 1d). Firstly, ST spots lying within regions containing cancer cells are indicated. Secondly, for each of such ST spots, we estimate the number of cells contained in that spot (using custom scripts in QuPath [25]; Methods). Next, we reconstruct cancer clones, their genotypes, and frequencies from the bulk DNA-seq data (using the existing methods Vardict [26], FalconX [27], and Canopy [9], see Methods, Fig. 1e). Afterwards, we analyse the data in the form of the number of alternated reads and the total number of reads for each mutation (mutation coverage), along with gene expression observed in each spot indicated as tumor (Fig. 1f). Notice, that the key assumption behind Tumoroscope is that each ST spot contains a hidden mixture of the clones reconstructed from the bulk DNA-seq data. Tumoroscope leverages: i) the estimated cell counts per spot provided as priors, ii) the alternated and total read counts for mutations in ST spots, and iii) the genotypes and frequencies for the clones using a probabilistic deconvolution model (Fig. 1g). The result of Tumoroscope is the identification of proportions of the clones in each spot (Fig. 1i). Additionally, for each spot, the method corrects the prior cell counts estimated from H&E images, using an inference from the ST data. Finally, we employ a regression model with gene expression data taken as independent variables and the inferred proportion of the clones in the ST spots as dependent variables (Fig. 1h) to infer gene expression profiles of the clonal populations (Fig. 1j).

**Fig. 1:**
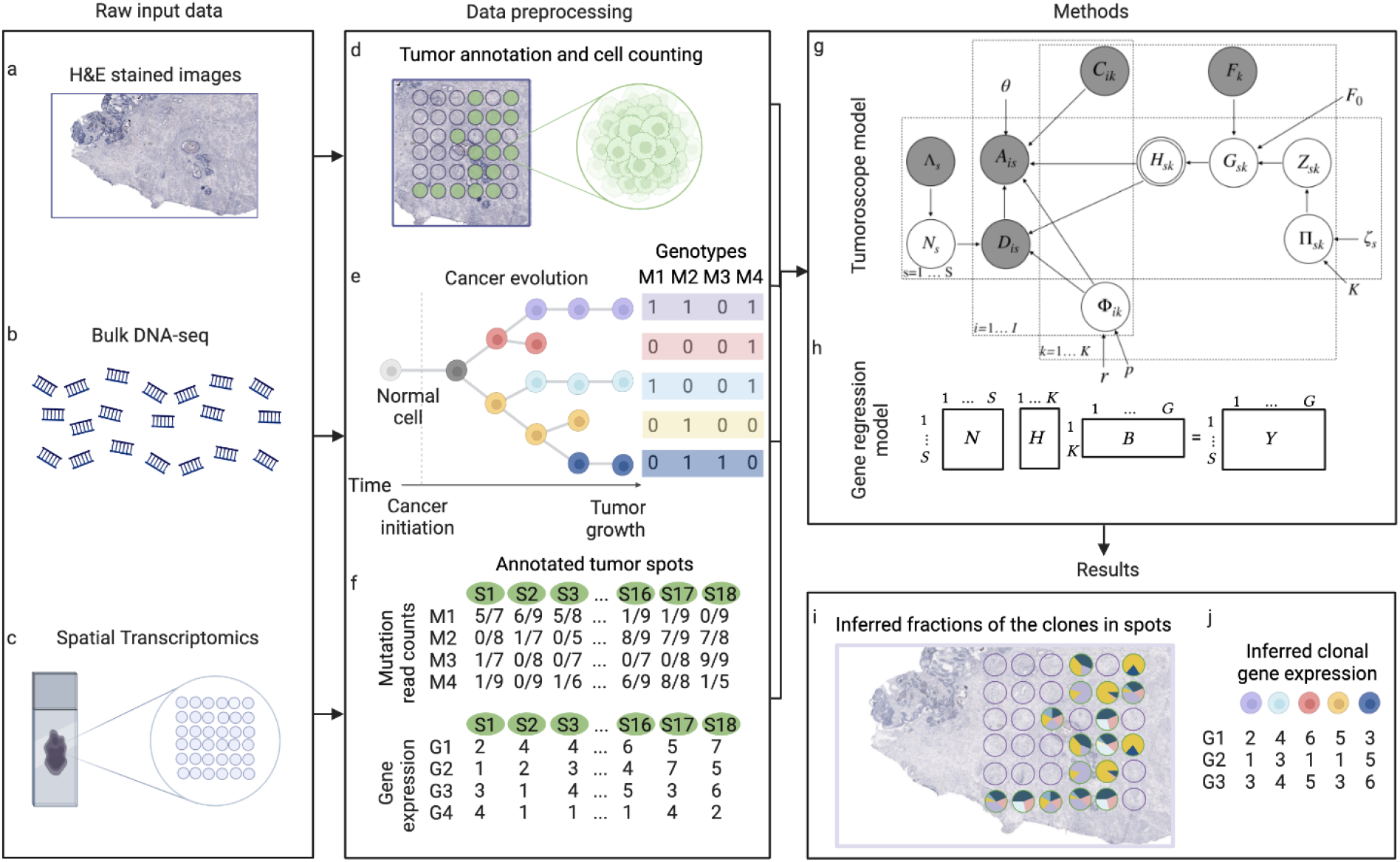
Tumoroscope framework overview. **a-c** Input data. **d-f** Data preprocessing. **g** Tumoroscope probabilistic model. **h** Regression model for inference of gene expression profiles of the clones. **i** Results of Tumoroscope. **j** Output of the regression model.

### Tumoroscope correctly estimates the proportion of clones in each spot and is robust to noise in input cell counts

In order to evaluate Tumoroscope’s performance in the case when the ground truth is known, we assessed its accuracy of estimating the proportion of clones in spots using simulated data. The simulation setups varied with respect to the number of mutations present in the clones, the expected number of clones in each spot, and the coverage of mutations. Specifically, we first designed a basic setup with five clones in the evolutionary tree, 30 mutations in the genotype matrix, an average number of 13.6 mutations per clone, and an expected number of 2.5 clones per spot. Next, we created four additional setups by decreasing and increasing the average number of mutations per clone to 5.1 and 15, respectively, and the expected number of clones per spot to 1 and 4.5, respectively. Furthermore, to test the influence of the level of coverage per mutation, for each of these five different setups we additionally varied the average coverage, with settings called very low, low, medium, and high (corresponding to an average number of reads present in each spot of 18, 50, 80, and 110, respectively; Methods). We simulated 10 datasets for each of the 20 setups resulting from the five aforementioned setups and four different coverage levels (amounting to 200 different simulated datasets used for evaluation in total; see Extended Data Table 1 for the detailed specification of simulation setups).

To inspect the robustness of our model to noise in the counts for number of cells per spot, we considered three different levels of noise in this input, as well as two versions of Tumoroscope, differing by how this input was modeled. Specifically, the model was either given the true simulated values of cells per each spot at the input, or we introduced small and large additive noise to these counts (Methods). In the first, default model version, referred to as Tumoroscope, the provided cell counts were used as priors and the number of cells per each spot was inferred accounting for all available data. In the second, simplified version, referred to as Tumoroscope-fixed, this input was used to fix the values of cell numbers in the spots. As both model variations were evaluated for the three levels of noise on each of the 200 simulated datasets, inference was made for 1200 synthetic datasets in total (Fig. 2). The performance was evaluated by calculating the Mean Average Error (MAE), that is, the average of the difference between the inferred proportions of the clone and the true values in all the spots and clones.

**Fig. 2:**
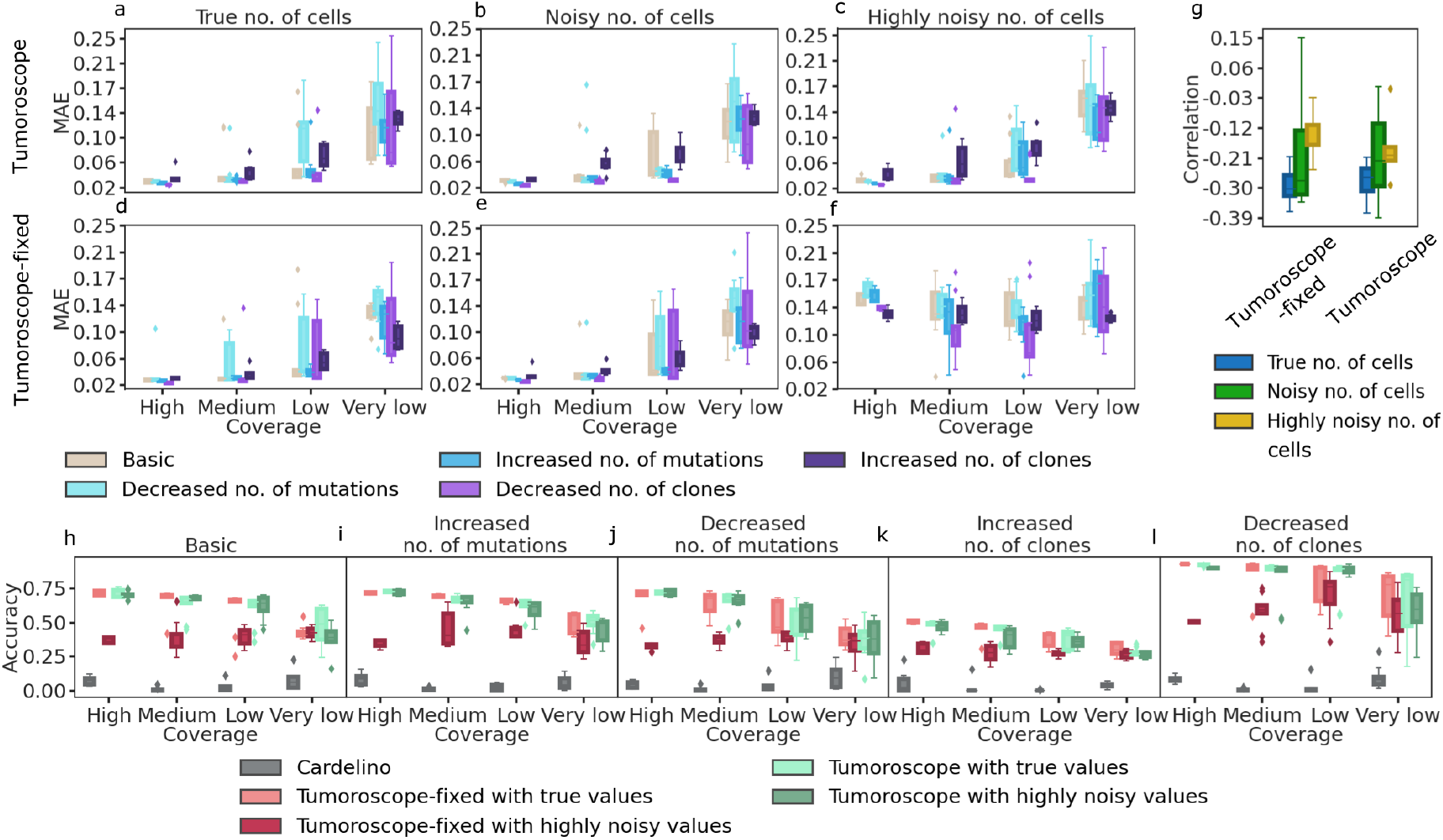
Performance of Tumoroscope on simulated data. **a-c** Mean Average Error (MAE; y-axis) as a function of mutation coverage (x-axis) in different simulation setups (colors) for Tumoroscope, for different noise levels in the cell count provided at input: no noise (**a**), medium noise (**b**) and high noise (**c**). **d-f** The same as in (**a-c**), but for Tumoroscope-fixed. **g** Correlation (y-axis) between the average mutation coverage and the average error in all the setups is negative for both model versions (x-axis), regardless of the noise in the number of cells provided at the input (colors). **h-l** Comparison of the accuracy (y-axis) of the model between cardelino (gray) and two versions of the model given true and highly noisy values for the number of cells (colors), depending on the mutation coverage (x-axis), in different simulation setups: basic (**h**), increased (**i**) and decreased (**j**) number of mutations, increased (**k**) and decreased (**l**) number of clones.

For both model versions, the error tended to increase with decreasing coverage (Fig. 2 a-g), indicating that better clone deconvolution can be obtained with deeper sequencing of the spots in ST data. Tumoroscope obtained low error (median MAE between 0.02 and 0.15, depending on the coverage), regardless of the level of noise in the input cell counts per spot (Fig. 2 a-c). Notably, in the case when the true, simulated cell counts were given as input, Tumoroscope performed equally well as Tumoroscope-fixed, despite the advantage that the latter had by fixing the counts to the true values (Fig.2a vs 2d). This advantage turned into bias when the input cell numbers became noisy, and Tumoroscope-fixed obtained a larger MAE than Tumoroscope (Fig. 2b,e, c, f). Similar results were obtained when higher coverages per mutation were considered (Extended Data Fig. 1). These results emphasize the importance of keeping the input cell count per spot as priors rather than fixing them as observed values, especially in the case of noise in this input, which is expected for real data. Indeed, in the real data, these input cell counts per spot are estimated from H&E images using algorithms for nuclei detection, which becomes particularly difficult when the cells are densely packed and the nuclei overlap (Methods).

### Accounting for the mixture of clones in each spot is key for model performance

To demonstrate the necessity of accounting for the mixture of clones in each spot, we compared Tumoroscope to an alternative method called cardelino [14]. Since cardelino was originally designed to assign single cells to clones based on scRNA-seq data, here, we applied cardelino providing each spot as a single cell, effectively assuming that the spot was a homogeneous readout from a single clone. For the sake of comparison, we only considered the major clone in each spot inferred by Tumoroscope and we defined the accuracy as the percentage of the agreement of the major inferred clone and the major true clone in the simulated data. Again, we evaluated both Tumoroscope and Tumoroscope-fixed. Tumoroscope obtained the worst-case median accuracy of around 0.27 for the increased number of clones and very low read count setup, and best-case accuracy of around 0.92 for the decreased number of clones and very high read count setup. With these results, Tumoroscope significantly outperformed cardelino, which obtained median accuracy between 0 and 0.09 in all simulation setups (Fig. 2 h–l). Similar to (Fig. 2 a–g), Tumoroscope’s accuracy tended to decrease with decreasing coverage. Interestingly, Tumoroscope obtained the highest accuracy for the decreased number of clones setup, and the worst for the increased nunmber of clones, indicating that the number of clones per spot is a decisive factor for method’s performance. Tumoroscope-fixed obtained lower accuracy than Tumoroscope, especially when provided with highly noisy input cell counts but still outperformed cardelino by a large margin. This result strongly emphasizes the importance of accounting for the clone mixtures in spots.

### Tumoroscope deconvolutes spatial clonal composition in a breast tumor and finds spatial patterns of cancer clones in sub-areas

To investigate the spatial clonal structure of a real tumor sample, we applied Tumoroscope to a newly generated dataset including three breast tumor sections from one patient. As input data, for each section, we generated deep whole-exome sequencing data (WES) and spatial transcriptomics (10x Genomics) of two neighboring layers (Methods). We assayed 4885-4992 spots per sample. In the data pre-processing stage, we selected the spots that were cancerous based on the expert pathologist’s annotations (Fig. 3e) and estimated cell counts from H&E images of the sections. We considered 608 high-confidence somatic single-nucleotide mutations (SNVs) identified from WES data that were co-observed in the annotated ST data (Methods). Next, we reconstructed the evolutionary tree of somatic mutations that were also present in ST data reads (Methods). We identified seven clones, including a base clone without somatic mutations (Fig. 3a,b; Extended data Fig. 3). Finally, given the selected 11461 cancerous spots, their estimated cell counts, total and alternated read counts at identified mutations, and the reconstructed clone genotypes, we used Tumoroscope to deconvolute the transcriptomics mutation profiles from the spots to obtain the proportions of the underlying clones.

**Fig. 3:**
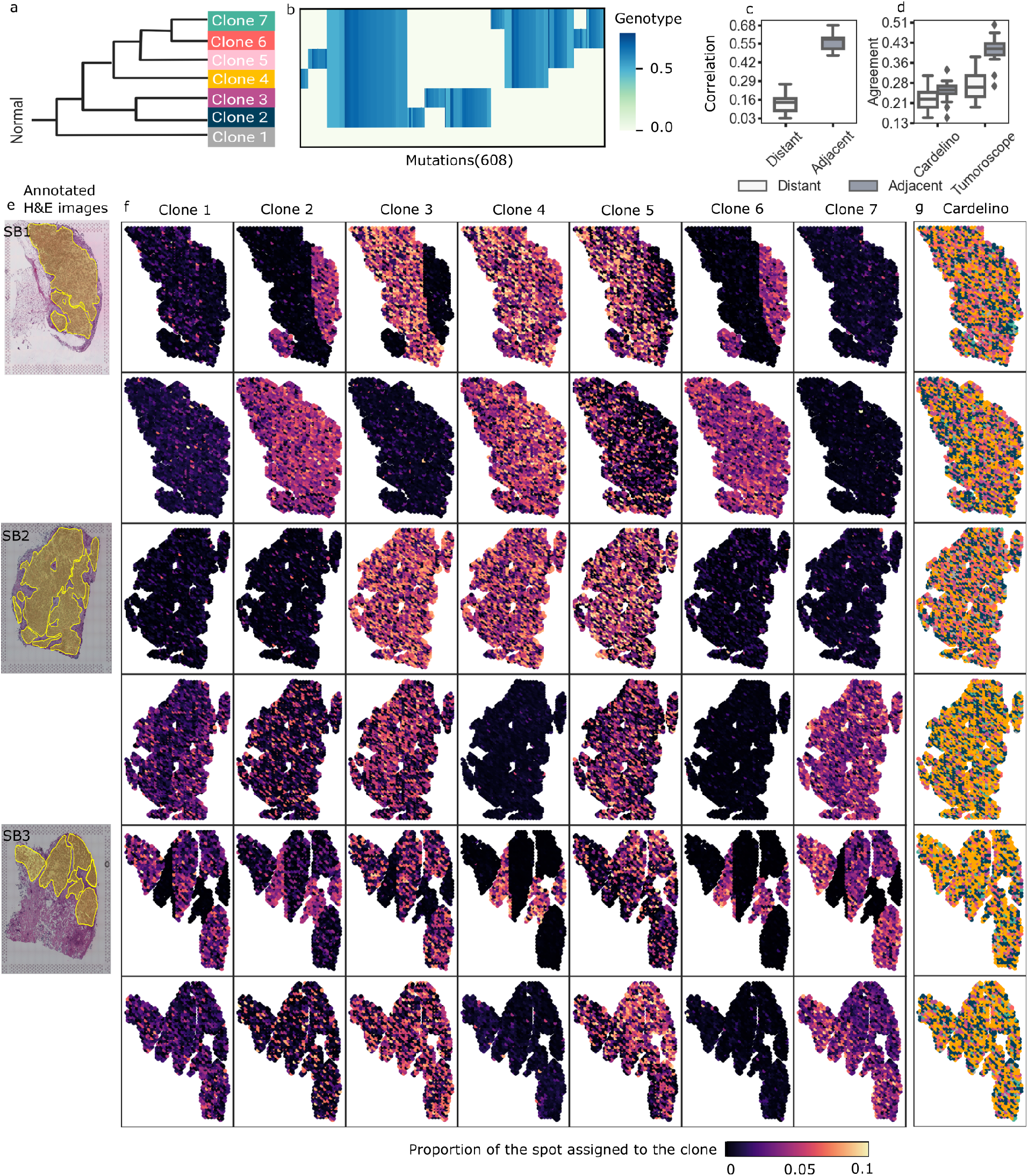
Spatial arrangement of cancer clones found for the breast cancer dataset. **a-b** Evolutionary tree and genotypes of the inferred clones. **c** Distribution of the correlation (y-axis) of the clonal composition of the spots that are distant and adjacent, computed for 100 pairs of spots sampled at random 20 times each (x-axis). **d** Distribution of the agreement of the distant and adjacent spots in cardelino and Tumoroscope, computed for the same randomly sampled pairs. For the computation of the agreement, we use the single inferred clone by cardelino and the major inferred clone by Tumoroscope. **e** Pathologist’s annotation of the cancerous areas on the H&E images for sections SB1, SB2, and SB3. **f** For each section, two rows correspond to the two nearby samples and 7 columns correspond to the proportion of the spots assigned to each clone. **g** The clonal assignment of the spots by cardelino for the same samples (see Extended data Fig. 2 for expanded cardelino results).

The composition of the seven clones in the investigated breast cancerous tissue identified by Tumoroscope revealed fascinating patterns of spatial arrangement (Fig. 3d). Generally, there was no single clone that fully dominated a specific contiguous sub-area of tissue. However, we did observe subsets of clones that coexisted in sub-areas. For section SB1, clone 4 was present in medium proportions in all analyzed spots of both layers. Very interestingly, there was a clearly separated sub-area in the right-hand part of section SB1 first layer, where clones 2, 4, and 6 co-occurred. The rest of this layer was dominated by clones 3, 4, and 5. In the second layer of section SB1, clones 2, 4, 5, and 6 coexisted, although with larger proportions of clone 4, low proportions of clones 2 and 6, and clone 5 being present in fewer spots than other clones. Clone 7 was not present in either layer of this section. Similarly, contiguous sub-areas that were predominantly occupied by small subsets of clones could be found in both layers of sections SB2 and SB3. As expected, clone 1, which lacked somatic mutations characteristic of the remaining cancerous clones, was found in only small proportions in the analyzed spots across all sections and layers. Patterns of clonal co-occurrence and mutual exclusion could be observed across all sections and layers, indicating a systematic mechanism. For example, the pairs of clones 2 and 6, as well as 3 and 5, although evolutionarily distant and with different genotypes (Fig. 3a, b) were always present together in the same sub-areas, while clones 4 and 7 excluded each other.

In contrast to Tumoroscope, in the assignments of single clones to spots inferred by cardelino, there was no detectable spatial pattern of domination of clones in sub-areas, as all clones were present in all sections uniformly (Extended Data Fig. 4 and 2). Again, this underlined the importance of spot deconvolution.

To validate the decomposition results in the absence of ground truth, we exploited that it is natural to expect the similarities in the clonal composition of the adjacent spots to be high, due to the growth process of the tumor in space. We found that the median correlation of clone proportions inferred by Tumoroscope between adjacent spots was significantly higher than the median correlation between distant spots (computed between 100 pairs of spots each, sampled at random 20 times; Fig. 3c). Since Tumoroscope treated each spot as independent and did not enforce any spatial similarities by design, this result strongly supports the correctness of the deconvolution of ST spots using Tumoroscope.

Importantly, we compared Tumoroscope’s and cardelino’s performances. Since cardelino was originally designed to analyze scRNA-seq data, when applied to ST data, it assigned only one clone to each spot. Thus, to enable the comparison, for every spot of interest, we determined the major clone (characterized by the highest proportion) indicated by Tumoroscope and computed the agreement for each out of 20 randomly sampled sets of adjacent and distant pairs of spots considered previously. The median fraction of adjacent pairs of spots with clonal assignment in agreement was much higher for Tumoroscope (0.41) than for cardelino (0.25). Moreover, the difference between the agreement for the distant and adjacent pairs was larger for Tumoroscope (distance between medians 0.14; one-sided Wilcoxon p-value 1.9e-06) than for cardelino (distance between medians 0.03; one-sided Wilcoxon p-value 0.063).

### Tumoroscope assigns the ST spots to clones in a prostate tumor

Next, we applied Tumoroscope to three prostate tumor sections from one patient, for which deep WES and ST data (custom arrays) of neighboring layers were generated [24]. As before, we selected the spots that were cancerous based on the regions that were annotated as tumor areas by an expert pathologist (obtaining 968-1001 spots per sample) and counted cells in spots from H&E images (obtaining 1-188 cells per spot; Fig. 4c; Methods). We then called the somatic mutations from WES data and identified 282 high-confidence somatic SNVs that co-occured in the ST data. Next, we reconstructed the evolutionary tree using Canopy [9] for that tumor from the WES data, identifying four clones, including a base clone without somatic mutations (Fig. 4a,b; Extended Data Fig. 4). Finally, we used Tumoroscope to deconvolute the transcriptomic signal from 294 spots in the ST data to reveal the proportions of the underlying clones.

**Fig. 4:**
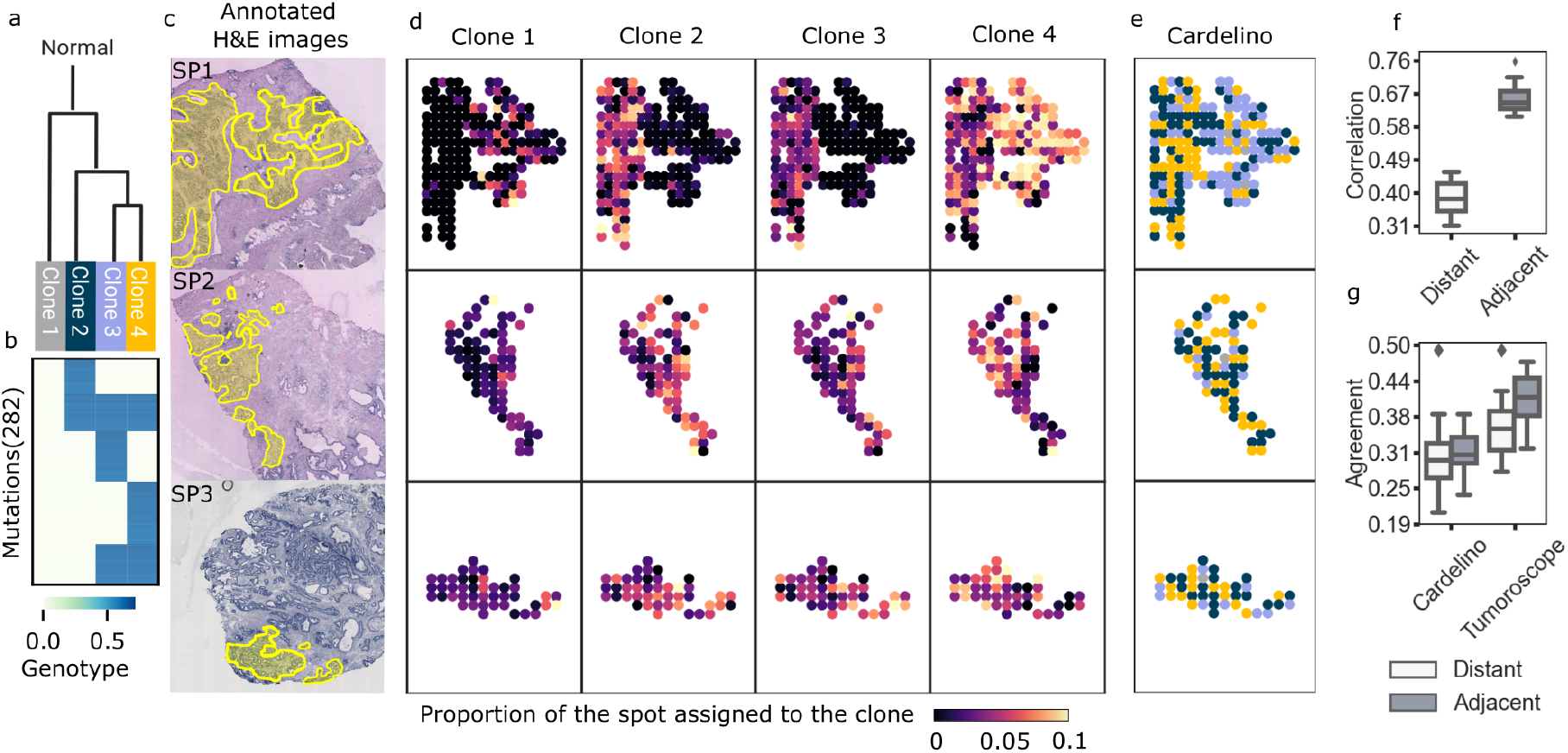
Results obtained for the prostate cancer dataset. **a-b** Evolutionary tree and genotype of the clones. **c** Pathologist’s annotation of the cancerous areas on the H&E images for sections SP1, SP2, and SP3. **d** For each section (rows), 4 columns correspond to the proportion of the spots assigned to each clone. **e** The clonal assignment by cardelino (see Extended data Fig. 5 for expanded cardelino results). **f** Distribution of the correlation of the clonal composition of the spots that are distant and adjacent, computed for 100 pairs of spots sampled at random 20 times each. **g** Distribution of the agreement (y-axis) of the distant and adjacent spots for cardelino and Tumoroscope, computed for the same randomly sampled pairs. For the computation of the agreement, we use the single inferred clone by cardelino and the major inferred clone by Tumoroscope.

Similar to the results obtained for breast cancer, for prostate cancer, we observed a pattern of sub-areas with marked presence of subsets of clones (Fig. 4d). Interestingly, section SP1 was divided into two sub-areas, with the left-hand sub-area containing all cancer clones 2, 3 and 4, while the right-hand sub-area was predominantly occupied by clone 4 with a small admixture of normal cells (clone 1). Sections SP2 and SP3 were smaller than SP1, but also showed distinct sub-areas with different clonal compositions.

For comparison, we again applied cardelino, by considering each spot in the ST data as a single cell measured using scRNA-seq (Fig. 4e). Interestingly, similarly to Tumoroscope, for section SP1 cardelino also divided the tissue into two different subareas, confirming their distinct clonal composition. However, the clones assigned by cardelino did not agree with the clones identified as taking the most proportion of the same spots by Tumoroscope. For example, for the right-hand sub-area of section SP1, cardelino mostly assigned spots to clone 3, and not 4.

We further verified whether Tumoroscope inferred more similar clonal profiles for adjacent spots than for distant spots. As expected, the correlations of the inferred clone proportions between of adjacent spots (median 0.65) were significantly higher than the correlations between distant spots (median 0.38; computed for 100 randomly selected pairs each and sampled 20 times; Fig. 4f), validating the results of Tumoroscope.

Furthermore, we compared the percentage of the agreement of the major clone in each spot in the adjacent and distant pairs of spots found using Tumoroscope, with the agreement of the clones in the same pairs of spots assigned by cardelino (Fig. 4g). With a median of 0.41, the agreement for adjacent spots was significantly higher for Tumoroscope than for cardelino (median 0.31). Furthermore, the difference between the agreement of the adjacent and distant spots was significant for Tumoroscope (difference between medians 0.05; one-sided Wilcoxon p-value 0.004) and was notably higher than for cardelino (0.01; one-sided Wilcoxon p-value 0.556).

### Similarity in gene expression profiles coincides with spatial co-occurrence of clones

Next, we applied the regression model (Methods) to deconvolve the expression values of genes in each clone from the aggregated gene expression values in spots. The regression model assumed that the expression of each gene in each spot is given by a mixture of expression values of that gene coming from clones present in that spot, weighted by their proportion inferred by Tumoroscope, and scaled by the inferred cell number in that spot. Both for breast and prostate cancer data, we ranked the genes by their maximum inferred expression across the clones and selected the first 30 genes at the top of the ranked list. We found that out of 30 genes selected for prostate cancer, 9 of them (*KLK2, KLK3, MSMB, TAGLN, SPON2, KLK4, PMEPA1, MYH11, AZGP1*) are known to be enriched in prostate cancer tissues (i.e, have elevated expression specifically in the prostate cancer tissue, according to Human Protein Atlas; HPA; [28]), and 19 are known to be upregulated in cancer (i.e, expressed in several cancer types according to HPA). For breast cancer, we found the *TPRG1* gene, known to be enriched in breast cancer tissues, and 25 genes known to be upregulated in cancer. The deconvolved expression profiles with respect to clones varied between the genes (Fig. 5). Using hierarchical clustering of the genes by their expression values, we found 10 clusters for breast, and 10 clusters for prostate cancer, respectively (Fig. 5a,b). Interestingly, we found clusters with genes expressed highly only in specific subsets of clones. For instance, for breast cancer data, we found that the genes *MT-CO1* and *MT-CO3*, associated with promoting cancer phenotype [29, 30] and gene *RPL19* associated with poor cancer patient survival [31], were highly expressed exclusively in clones 2, 6 and 7. Furthermore, in the results obtained for prostate cancer data, we found that the gene *KLK3*, known as prostate-specific antigen and the most frequently clinically used prostate cancer biomarker [32], was active in clones 2 and 3. These results suggest that each cancer cell clone might have its specific function in tumor progression and development.

**Fig. 5:**
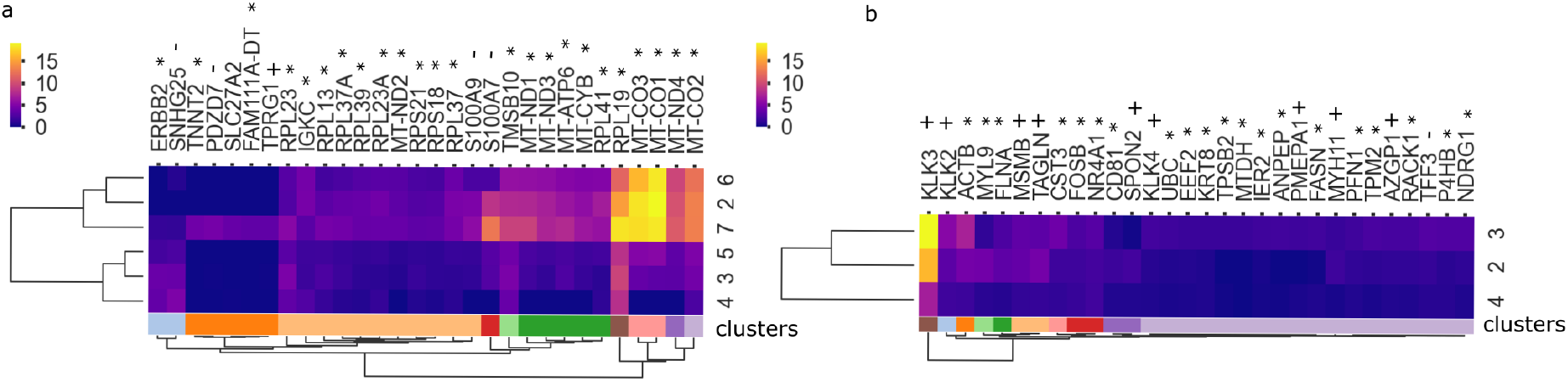
Genes are expressed differently in various cancer clones. The expression of the 30 genes that were inferred by the regression model as the most active in at least one clone, clustered in rows and columns, for breast (**a**) and prostate cancer (**b**) tissues. * cancer gene found in all cancer tissues (not cancer type specific) according to the HPA database [28]; + cancer gene with nTPM (normalized gene expression value) in the desired cancer type (either breast in **a** or prostate in **b**) at least four times higher than in other cancer tissues, according to [28]; ++ cancer gene with nTPM in a group of cancer tissues including the desired cancer type, at least four times higher than in other cancer tissues, according to [28]; - not detected in cancer tissues, or nTPM at least four times higher in another cancer tissue than the desired one, according to [28].

Finally, we performed clustering of the clones with respect to their gene expression profiles (Fig. 5). Intriguingly, for both prostate and breast cancer, clones with similar inferred phenol-types, which were clustered together by their expression profiles, also coexisted in space across the tissue (compare with Fig. 3 and 4). For breast cancer, since the correlation between the fraction of the clones 2 and 6 in the spots was relatively high (Pearson correlation *r* = 0.64; see Extended data Fig. 7), it was expected to find them to be similar in terms of their gene expression profiles, per construction of the regression model. In contrast, while the fractions of clones 3 and 5 in the same spots were not correlated (Pearson correlation *r* = −0.28; Extended data Fig. 7), they were still co-localized in tissue space as they co-occurred in adjacent spots (Fig. 3; average correlation of fractions in the adjacent spots *r* = 0.16; Extended data Fig. 8). In this case, similar expression profiles for these clones were not expected per construction of the regression model, but still, these clones were inferred as the second most similar. For the prostate cancer sample, the two clones 2 and 3 which were co-localized in the tissue (Fig. 4), were also the most similar in terms of their gene expression profiles. Both for breast and prostate cancer, the pairs of correlated clones were not close in terms of their mutations (Fig. 3b) and thus placed in distinct sub-trees of the evolutionary tree (Fig. 3a). This indicates that even distant clones may have similar phenotypes and play analogous roles in tumor progression.

## Discussion

Tumoroscope is the first approach for mapping cancer clones based on point mutations in tissue space and resolving their expression profiles in high resolution. This resolution amounts to the diameter of the deconvoluted spots, ranging from 100 µm (as for the prostate cancer dataset [24]) to 55 µm (as for the breast cancer dataset), depending on the ST technology. Effectively, this means the model is able to assign clone proportions for spatially resolved mini bulks of the order of 1-40 cells (1-10 cells for breast cancer dataset [23] and 10-40 for prostate cancer dataset [22]). Tumoroscope achieves this result by innovative integration of data from technologies that were not originally developed for this task: H&E, WES, and ST. The key signal exploited by Tumoroscope to identify the clonal composition of ST spots is the matching between mutations present in the genotypes of the clones to the mutations found in the RNA sequencing of the spots. On top of that, the method estimates additional variables, such as the number of cells in each spot and the average expression of each variant site per single cell. Finally, with the proportion of the spots coming from specific clones, alongside the gene expression observed in spots in hand, we solve the problem of clone-specific gene expression deconvolution.

Our comprehensive simulation study demonstrates Tumoroscope’s robustness to noise in the estimation of the number of cells in ST spots. The results clearly indicate that the deconvolution task becomes easier with increasing coverage of mutations in ST spots and with a decreasing number of coexisting clones in each spot. In application to breast and prostate cancer data, Tumoroscope reveals spatial patterns of clonal arrangement, indicating a well-mixed coexistence of small subsets of all clones in subareas of the tumor tissue.

Applying our regression model to infer gene expression levels in the different clones allows us to identify the distinct phenotypes of the clones, effectively assigning spatial resolution to the function of the different tumor subpopulations, and thus profiling the functional heterogeneity of tumors. Moreover, our findings in both analyzed cancer types indicate that it is the phenotypic, and not genotypic similarity, which could drive the spatial co-occurrence of clones. However, this result should be further validated in additional patient samples and using independent data.

To our knowledge, there exists no competing technology that could be applied to resolve the spatial clonal heterogeneity of tumors in a comparable resolution to Tumoroscope. Spatial capturing of DNA sequences is still at the very early stage of development [33]. The very low resolution obtained with current spatial DNA sequencing technology requires merging beads located nearby in the array and, thus, provides spatial mini-bulk data, akin to ST spots. Additionally, ignoring the evolutionary origin of distinct clones and clustering beads with no information about variant allele frequency, as performed in [33], oversimplifies the complex problem of spatial clonal deconvolution. Considering the difficulties intrinsic to spatial DNA-seq data, high-resolution ST data proves to be a highly attractive alternative for the spatial inference of clonal evolution. The recently developed method STARCH [34] combines RNA-sequencing of ST spots and DNA-sequencing from neighboring tissues in the same tumor sample to infer the spatial arrangement of clones based on their copy number profiles. Besides, Erickson *et al*. [35] developed a method to infer genome-wide copy number variations (CNVs) from spatially resolved mRNA profiles in situ that reveals distinct CNV based clonal patterns within tumors. In contrast to Tumoroscope, however, both these methods do not directly address the problem of deconvoluting the mixture of cancer clones per each spot.

The quality of the obtained results could be further improved with better technology. For example, replacing WES with scDNA-seq data would allow more accurate inference of cancer clones, their evolutionary relationships, and genotypes using dedicated computational approaches [36, 37]. As the ST technology improves, smaller spots are expected, limiting the number of clones per spot, which makes the deconvolution problem easier. Finally, currently, only the first 300 bp of gene sequences are sequenced in the process of ST data generation. For our approach, ideally, whole gene bodies should be sequenced so that all mutations detectable from WES could also be observed in ST and matched for more accurate deconvolution of spots into clones. Such a sequencing was recently shown to be possible [38] but was not available for the data that we analyzed and is not in the standard ST protocols.

Despite these technological limitations, already now Tumoroscope offers a major break-through in the integrated analysis of spatial, genomic and phenotypic tumor heterogeneity. The model could be applied in further studies profiling adjacent tumor samples to provide 3D maps of clones. With our ability to compute gene expression profiles of the clones, we could make it possible to predict the most proliferating areas and, thereby, the most probable expansion sites of the 3D structure of the tumor. Furthermore, studies combining H&E, WES, and ST for large cohorts of patients, could explore the dependencies between patient clinical features and the spatial patterns of clones found using Tumoroscope. Combined with cell-type deconvolution approaches for ST data in the tissue surrounding the tumors [39–41], our framework has the potential to bring unprecedented insights into the interactions of specific cancer clones, their phenotypes, and the surrounding microenvironment. In summary, Tumoroscope opens up a new avenue in cancer research with broad applications for a basic understanding of the disease and its clinical applications.

## Methods

### Breast tumor samples

The breast tumor study was approved by the Swedish Ethical Review authority (no. 2016/957-31 with amendments 2017-742-32, 2020-00323 and 2021-00795). Breast tumor tissues were obtained by Dr. Johan Hartman (Institute of Oncology and Pathology, Karolinska Institute). The samples were collected from tumor material removed from a patient with untreated invasive ductal carcinoma during breast cancer surgery. Histological evaluation of the patient’s tumor was performed by pathologists for diagnostic purposes and defined as HER2 (+3), ER (30%), and Ki67 (79%). For this tumor sample, different regions (*n* = 5) were selected by the pathologists. From each region, tissue was isolated for immediate embedding in OCT for gene expression analysis with spatial transcriptomics. Samples for spatial transcriptomics were immediately frozen and stored at −80°C until further analysis.

### Preparation and sequencing of Spatial Gene Expression Libraries for the breast tumor samples

Sections of fresh-frozen breast tumor tissue were cut at 10 µm thickness and mounted onto slides from the Visium Spatial Gene Expression Slide & Reagent kit (10X Genomics). Sequencing libraries were prepared following the manufacturer’s protocol (Document number CG000239 Rev A, 10x Genomics). Prior to imaging, coverslips were mounted on the slides according to the protocol’s optional step Coverslip Application & Removal. Tissue images were taken at 20x magnification using Metafer Slide Scanning platform (MetaSystems) and raw images stitched with VSlide software (MetaSystems). Adaptations of the protocol were made in that the Hematoxylin staining time was reduced to 4 minutes and tissue permeabilization was performed for 12 minutes. Final libraries were sequenced on NextSeq2000 (Illumina) or NovaSeq6000 (Illumina).

### Data processing of spatial gene expression libraries for the breast tumor samples

Following demultiplexing of bcl files, read 2 fastq files were trimmed using Cutadapt [42] to remove full-length or truncated template switch oligo (TSO) sequences from the 5’ end (beginning of read 2) and polyA homopolymers from the 3’ end (end of read 2). The TSO sequence (AAGCAGTGGTATCAACGCAGAGTACATGGG) was used as a non-internal 5’ adapter with a minimum overlap of 5, meaning that partial matches (up to 5 base pairs) or intact TSO sequences were removed from the 5’ end. The error tolerance was set to 0.1 for the TSO trimming to allow for a maximum of 3 errors. For the 3’ end homopolymer trimming, a sequence of 10 As was used as a regular 3’ adapter to remove potential polyA tail products regardless of its position in the read, also with a minimum overlap of 5 base pairs. The trimmed data was processed with the spaceranger pipeline (10X Genomics), version 1.0.0 (BC) and mapped to the GRCH38 v93 genome assembly.

### Prostate cancer sample

The prostate cancer dataset was generated and published by Berglund *et al*. [24]. This dataset consists of twelve sections, with H&E images, bulk DNA-seq and spatial transcriptomics provided for each section. The data were generated and processed using protocols as described in [24].

### Identifying the spots that contain tumor cells

To select the spots that contain tumor cells, we took advantage of H&E staining images of the analyzed tissues. For both breast and prostate cancer, regions containing cancer cells were annotated by an expert pathologist Dr. Łukasz Koperski using QuPath [25]. We further selected spots whose area overlapped with the pathologist’s annotated regions, using a custom script in QuPath [25].

### Counting cells in spots

We developed a custom script in QuPath [25] to count cells in each ST spot visible in the H&E images [25]. The script takes as input coordinates and diameters of spots to define target areas. Then, we employ QuPath’s inbuilt cell counting algorithm for detecting and counting nuclei. In order to adjust parameters of the algorithm, we examined random spots by manually counting cells to verify the accuracy of the results.

### Spatial transcriptomics data preprocessing

For prostate cancer sample, the ST data bam files were provided by Berglund *et al*. [24]. For breast cancer sample, to create the genome index, we used the STAR program [43] with the GRCh38 reference genome as input. Next, we applied the ST Pipeline [44], providing the genome index, FASTQ files, barcodes and array coordinates as input. We obtained the gene expression matrix as counts of reads for each gene, which the ST Pipeline produces by default. In addition, we modified the default settings, to obtain BAM files with the mapped reads.

### Bulk DNA-seq and somatic mutation calling

We identified somatic mutations that appeared in at least one of the bulk DNA-seq sections, by calling the mutations using Vardict [26] for each section with a p-value threshold equal to 0.1. Then we used their union over sections as the set of mutations called in bulk DNA-seq data. This procedure was performed in the same way for the prostate and the breast dataset.

### Selection of somatic mutations that are detected both in bulk DNA-seq and ST data

Next, we identified the bulk DNA-seq mutations that were also present in ST data. For calculating the total and alternated reads over the mutations in ST data, we located the selected bulk DNA-seq mutations in the ST bam files and counted the corresponding mapped reads with our script. The reads with a different nucleotide as compared to the reference genome were called the alternated reads.

Finally, we selected the mutations for which there existed at least one alternated read in at least one section. The alternated and total read counts in bulk DNA-seq data for the selected mutations were given as input for phylogenetic inference, while the alternated and total read counts in ST data for the same mutations were given as input to Tumoroscope. The median read coverage for the selected variant sites in bulk DNA-seq data for breast and prostate cancer were 214.5 and 134.75, respectively.

### Phylogenetic tree analysis

To identify the phylogenetic tree and infer the genotype and prevalence of each clone in the tree we used a statistical method called Canopy [9]. The input to Canopy are variant allele frequencies of somatic single nucleotide alterations (SNAs), along with allele-specific coverage ratios between the tumor and matched normal sample for somatic copy number alterations (CNAs). We used FalconX for producing the allele-specific coverage ratio between tumor and normal sample [27]. We used multi-sample feature of Canopy to infer the clonal evolution across the sections for both prostate and breast datasets.

### Mapping fractions of cells in ST spots to cancer clones using Tumoroscope

Tumoroscope is a probabilistic graphical model for estimating proportions of cancer clones in ST spots given alternated and total read counts over the analysed somatic mutations, genotypes and frequencies of the clones, and estimated cell counts per each spot (Fig. 1f). Let *i* ∈ {1, …, *M*} index the selected mutation positions, identified both in bulk DNA sequencing and ST data. We are given a set of *K* cancer clones, indexed by *k* ∈ {1, …, *K*} as input, which has been derived from bulk DNA sequencing data. The genotypes of the input clones are represented as a matrix *C* with entries between 0 and 1 corresponding to the zygosity. *C*_*i,k*_ equals 0 if there is no mutation on position *i* in clone *k*, equals 1 in case all alleles of that position carry the mutation, and equals 0.5 when the half of the alleles of that position carry the mutation. Note that there can be multiple alleles for position *i*. In general, the zygosity is defined as the ratio of the number of mutated alleles to the total number of alleles and we estimate it by the ratio of the major allele frequency to the total read count. The prevalence of the clones in the bulk DNA sequencing is represented by the vector *F* = (*F*_1_, …, *F*_*K*_), with values summing up to one. Let *s* ∈ {1, …, *S*} index the spots. We use a feature allocation model to account for the presence of clones in spots [45]. Specifically, we define *Z*_*s,k*_ ∈ {0, 1} as an indicator of the presence of clone *k* in spot *s*. We assume a Bernoulli distribution over *Z*_*s,k*_ and a Beta prior over its parameter Π with hyper-parameter *ζ*_*s*_:

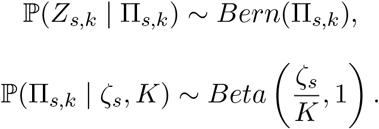

Let **1** = {1}^*K*^ denotes a *K*-dimensional vector with all elements equal to 1. Bearing in mind the assumption about Beta prior over Π_*s,k*_, we calculate the expected number of nonzero entries in each spot 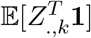 using the formula for the mean of the Beta distribution as [46, 47]

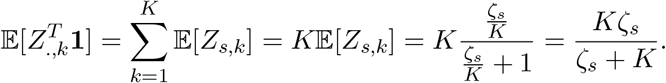

Given this formula and the number of the clones *K*, we are able to control the expected number of clones in each spot by tuning shape parameter of the beta distribution, 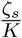.

Our main goal is to estimate the proportions of clones in the spots, which are represented by the variable *H*, a matrix with *S* rows and *K* columns. The value of an element *H*_*s,k*_ is the fraction of spot *s* coming from clone *k*. We consider a Dirichlet distribution over *H*_*s*,._ = (*H*_*s*,1_, …, *H*_*s,K*_),

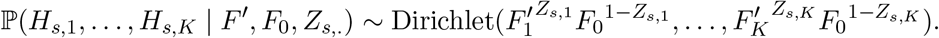

Here, *F*_0_ corresponds to a “pseudo-frequency”, and results in non-zero proportions for all clones for each spot. We set *F*_0_ to a small number, effectively assigning small proportions to clones which are not present in the spot. The 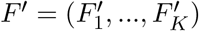 are obtained as discretized frequencies *F*. Specifically, we discretize the values of *F* by dividing the range from 0 to 1 into 20 equal-sized bins and then round up the values to the upper-bounds of the bins and scale them by multiplicative factor *l*

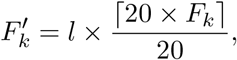

where we used *l* = 100, but it can be specified by the user.

To sample *H*, we take advantage of the relation between Dirichlet and Gamma distribution [48] and draw *K* independent random samples (*G*_*s*,1_, …, *G*_*s,K*_) from *K* Gamma distributions,

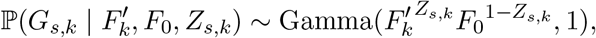

and then we calculate the proportions *H*:

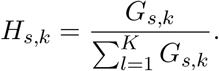

The total read count at position *i* in spot *s* is represented by observed variable *D*_*i,s*_. We assume a Poisson distribution over *D*_*i,s*_,

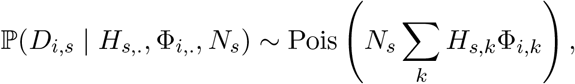

where Φ_*i,k*_ is the average coverage for the position *i* across the cells from clone *k*, and *N*_*s*_ is the number of cells in spot *s*. The variables *N*_*s*_ can be fixed to *a priori* known values.

However, in most practical applications, the number of cells per spot is not known. This gives a compelling reason to estimate them as a part of model inference. We assume a Poisson distribution over *N*_*s*_,

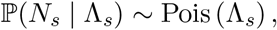

where Λ_*s*_ is the expected number of cells in spot *s*. Also, we assume a Gamma distribution over Φ_i,k_,

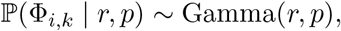

where *r* and *p* are the shape and rate hyperparameters, respectively.

*A*_*i,s*_ represents the number of alternated reads for position *i* in spot *s*. We assume a Binomial distribution over *A*_*i,s*_,

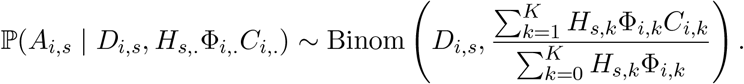

Where the success probability of Binomial distribution is the probability of observing *A*_*i,s*_ alternated reads out of *D*_*i,s*_ reads in total. Given the variables *N*_*s*_,*H*_*s*,._ and Φ_*i*,._,we calculate the expected number of alternated reads and the total reads in spot *s* using 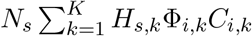 and 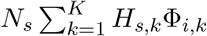 respectively. Therefore, we calculate the success probability by calculating the fraction of the expected number of alternated reads and the total reads.

### Metropolis-Hasting inside Gibbs Sampling

In the Gibbs sampling, we iteratively generate samples from each hidden variable’s conditional distribution, given the remaining variables, in order to estimate the posterior distribution of the hidden variables. Each hidden variable given the variables in its Markov Blanket is conditionally independent of all variables outside its Markov Blanket in the graphical model [49]. A variable’s Markov Blanket includes its parents, children, and children’s parents. If the conditional distribution does not have a closed analytical form, we use a Metropolis-Hasting step inside the Gibbs sampler. In the following, we describe the sampling steps for each hidden variable.

### The variables with the closed-form sampling distribution

Π_*s,k*_ and *Z*_*s,k*_ are the only variables with analytical sampling distributions.

### Sampling Π_*s,k*_

For sampling Π_*s,k*_ we take advantage of the conjugacy of Beta and Bernoulli distributions:

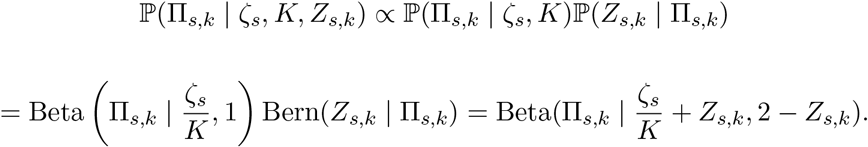

### Sampling *Z*_*s,k*_

For sampling *Z*_*s,k*_ we utilize the fact that this variable only accepts binary values. Therefore, we sample 0 or 1, proportional to their corresponding calculated probabilities.

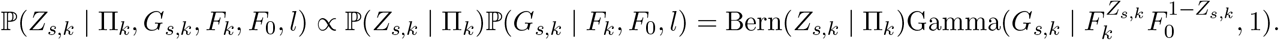

### Metropolis-Hasting adaptive steps inside Gibbs sampler

In our model, there is no closed analytical form of conditional distribution for variables Φ_*i,k*_, *G*_*s,k*_ and *N*_*s*_. Therefore, we take advantage of Metropolis-Hasting inside Gibbs sampler. We compute the acceptance ratio *A* as the following

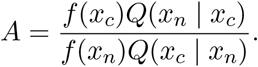

Where *f* (*x*) is a function that is proportional to the desired density function *P* (*x*) and *Q* is the proposal distribution. Bearing in mind the non-negativity of the variables of our interest, we choose a Truncated Normal distribution for *Q* with the mean value of the current sample *x*_*c*_ and variances *σ*_Φ_, *σ*_*G*_ and *σ*_*N*_ corresponding to each variable. The variance of the Truncated Normal distribution determines the proximity of the new sample from the current one, which is interpreted as the step size. The choice of the step size has a major impact on the acceptance rate of the Metropolis Hasting. We tune the *σ*_Φ_, *σ*_*G*_ and *σ*_*N*_ every *b* steps starting with an arbitrary value based on the feedback from the acceptance rate. Firstly, we choose an optimal acceptance rate *R*_*o*_ for each variable. Secondly, we modify the variance by *δ* percent of the current variance and *δ* is calculated by the difference of the optimal and current acceptance rate *R*_*c*_. Ultimately, during the sampling steps, we learn the optimal variance value for each variable.

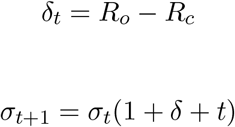

In the following, we describe the conditional distribution for each variable.

### Conditional distribution for Φ_*i,k*_

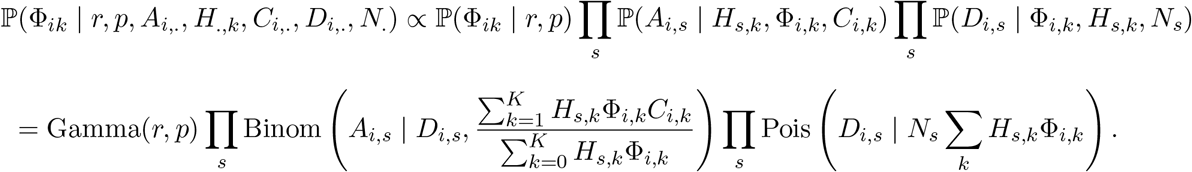

### Conditional distribution for *G*_*s,k*_

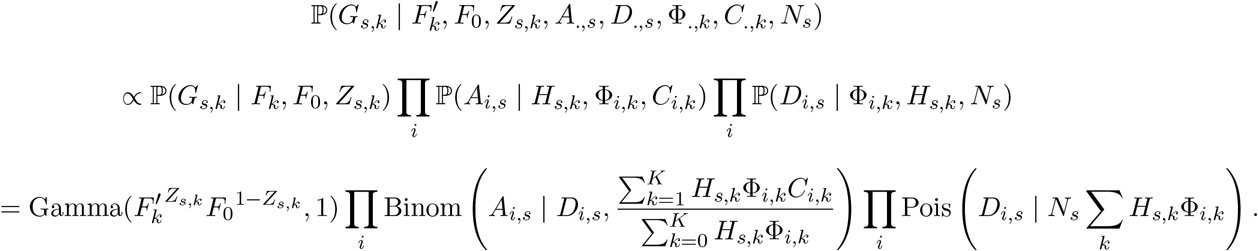

### Sampling *N*_*s*_

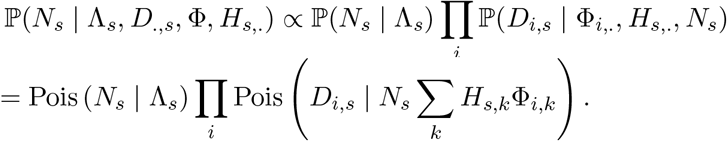

### Parameter setting for different simulation setups

First, we calculate the parameter of the Beta distribution over variable Π_*s,k*_ based on the assumed expected value of the number of clones:

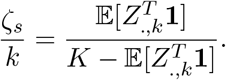

Considering expected values of 1, 2.5, and 4.5 for the number of clones found in each spot, we obtain 0.25, 1, and 9 and use these values for the Beta distribution parameter.

Second, we exploit Φ_*i,k*_ that represents the expected number of reads for mutation *i* in each cell for generating different read coverage for total and alternated read counts. We set *p* = 1. With this, we control the expected value of Φ_*i,k*_ using parameter *r*.

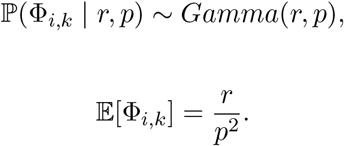

For the very low, low, medium and high number of reads, we consider *r* = 0.02, *r* = 0.07, *r* = 0.09 and *r* = 0.19, respectively, leading to the 18, 50, 80, and 110 average total reads for each spot.

Last, we generate three datasets for the number of cells with different level of noise to compare our two models having number of cells as observed and hidden variable. We add the noise value *ϵ* to the true values.

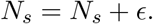

We consider *ϵ* = 0, *ϵ* ∼ Pois(1) and *ϵ* ∼ Pois(10) for generating without noise, noisy and highly noisy number of cells.

### Parameter estimation obtained for the real data

For the higher accuracy of the graphical model reflecting the real data, we estimate the input parameters of the model based on the characteristics of the data. The first parameter is *λ*_*s*_, the expected number of the cells in spot *s*, which affects the estimation of the number of cells and, ultimately, the number of reads we are expecting, which is a crucial element for estimating the fraction of the clones. Therefore, we estimate the number of cells using the H&E images and a customized script in QuPath and use them as the mean parameter for the Poisson distribution over *N* (described above) [25]. Next parameters are *r* and *p*, the shape and rate in the Gamma distribution over variable Φ. We use mixed type log-moment estimators for calculating *r* and *p* [50].

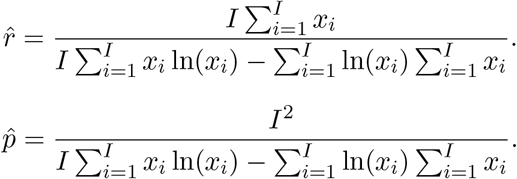

Where *x*_*i*_ with *i* ∈ {1, …, *I*} are the sample from Gamma distribution. We generate these samples using the total number of reads *D*. We calculate the average number of reads from every cell dividing the reads from the spots to the number of estimated cells as input which gives us *I* samples, equal to the number of mutations.

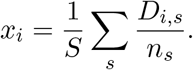

### Clonal composition resemblance in adjacent spots

The evolutionary process imposes the similarity of the clonal composition in the adjacent spots. Therefore, we expect to have a higher correlation between the clonal composition of the adjacent spots as compared to distant spots. To make this comparison, we randomly generate *N* pair of adjacent spots 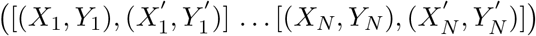 with *X* and *Y* corresponding to their coordinates. These adjacent pairs satisfy two constraints of 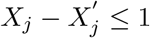 and 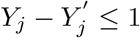 indexed by *j* ∈ {1, …, *N*}. We also generate *N* pair of distant spots with the two constraints of 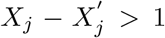 and 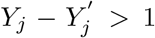 We define 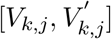 as the fraction of clone *k* in spots corresponding to the *j*^*th*^ pair in the adjacent spots. Then we calculate the Pearson correlation for the vector 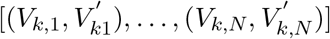. The procedure is repeated for all the clones and distant spots for the sake of comparison.

### Clonal assignment of the spots using cardelino

Cardelino [14] is a statistical method originally developed for inferring the clone of origin of individual cells using single-cell RNA-seq (scRNA-seq). It integrates information from imperfect clonal trees inferred from whole-exome sequencing data and sparse variant alleles expressed in scRNA-seq data. However, here, we applied it on spatial transcriptomics instead of scRNA-seq to validate the assumption of mixture of clones in each ST spot instead of assuming homogenous spots contaning only one clone. We used *clone*_*id* function with “sampling” inference mode, minimum iteration of 100000 and maximum iteration of 250000. We used 3 parallel chains for prostate cancer data and 1 chain for breast cancer data due to the high RAM demand of the cardelino.

### Estimating gene expression of the clones

Having the proportions of the clones in each spot inferred using Tumoroscope and gene expression data from spatial transcriptomics, we estimate average clonal gene expression using a regression model. Let *g* ∈ {1, …, *G*} index genes and *Y* be a matrix with *S* rows and *G* columns, where *Y*_*s,g*_ is the measured gene expression of gene *g* in spot *s*. We are interested in estimating *B*_*k,g*_ - average gene expression of gene *g* in one cell of clone *k*. We use *H* and *N* variables inferred by Tumoroscope, and we rewrite *N* as an *S* × *S* diagonal matrix *N′*, where 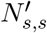 is the number of cells in spot *s* and other elements of the matrix are equal to zero. We describe the relationship between the variables with an overdetermined system of equations *N′HB* = *Y*. Then we try to find the optimal solution of this equation using linear regression with a lower bound of *B*_*k,g*_ ≥ 0 and no intercept. For this purpose, we apply a python function *scipy*.*optimize*.*lsq*_*linear* to the data.

## Data and Code availability

Tumoroscope can be obtained as an installable Python package, via ‘pip install tumoroscope’, and is available under the GNU General Public License v3.0. Tumoroscope implementation, package updates, and datasets supporting the conclusions of this article will be maintained at https://github.com/szczurek-lab/Tumoroscope.

## Acknowledgements

We acknowledge Andrew Roth for suggesting the feature allocation model as part of Tumoroscope.

## Author contributions

S.D.S., E.S., J.L. and A.G. developed the probabilistic model. S.D.S. implemented the model and carried out the application of the model, supervised by E.S. B.J. applied the regression on gene expression data and performed GSEA analysis. J.H. extracted the breast cancer tumor. X.C. prepared the sample, performed the DNA extraction for whole-exome sequencing of breast cancer dataset. K.T. and C.E. performed spatial transcriptomics. J.M. and C.E. performed the cell sorting. A.S.N. performed st pipeline and bulk DNA-seq mutation calling. S.D.S., J.L. and E.S. conceived the study. S.D.S., E.S., A.C., A.G. and J.L. wrote the paper. S.D.S. carried out the bench-marking of alternative methods. Ł.K., Ł.R and I.F. analysed the H&E images. S.D.S., A.C., D.N., Ł.K. analysed and interpreted the model results. All authors provided critical feedback; helped shape the research and analysis; edited, reviewed and approved the manuscript.

## Extended data

**Extended data Figure 1:**
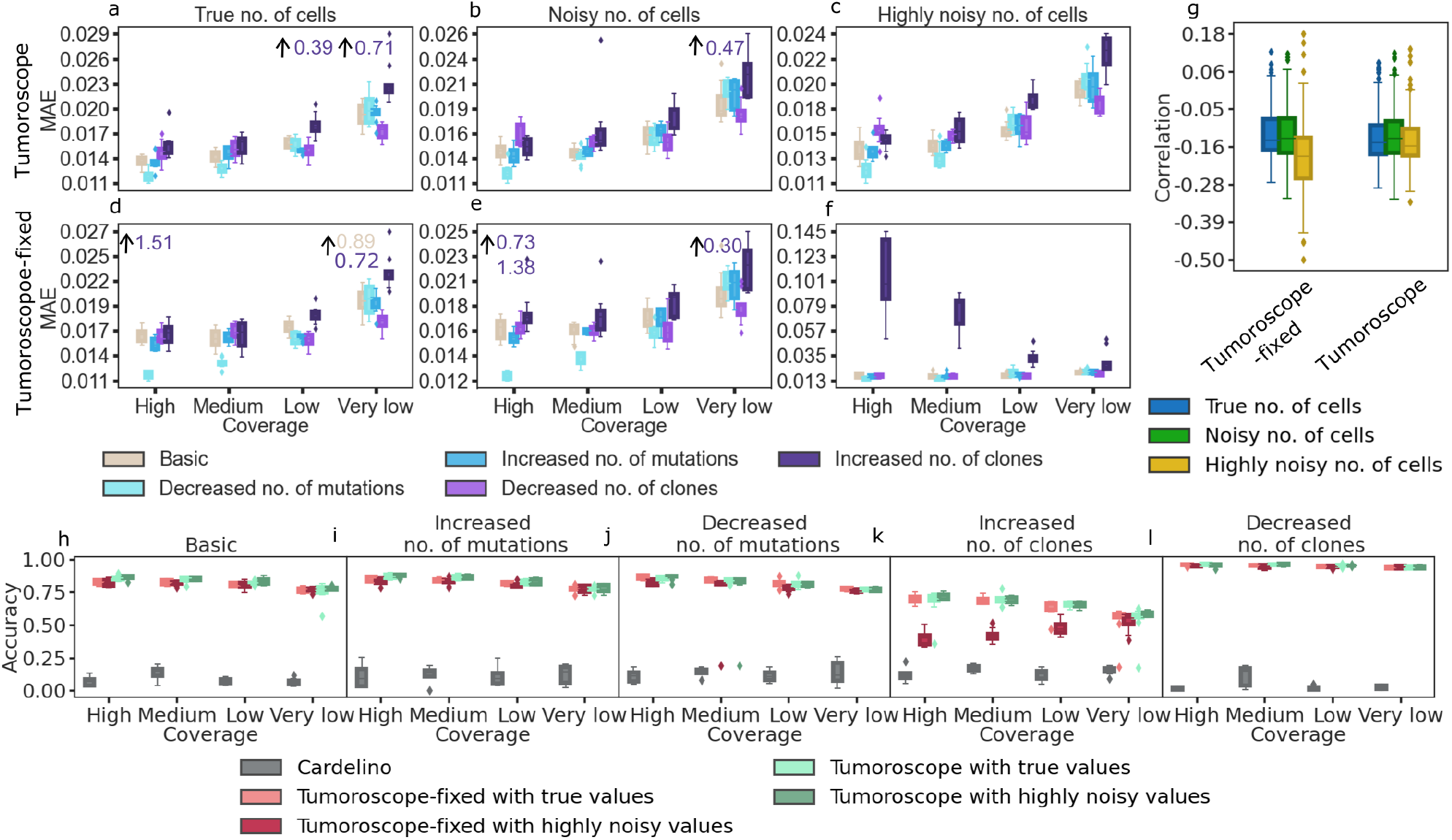
Performance of Tumoroscope on simulated data with a high number of reads. The axess and legends are the same as in Fig. 2 in the main text. Here, high, medium, low, and very low are corresponding to the average number of reads present in each spot of 297, 734, 1488, and 2246, respectively. Overall, compared to Fig. 2, having a higher number of reads increased the performance (strongly decreased MAE) for the estimation of the fraction of the clones in spots. **f** In the case when Tumoroscope-fixed is given a fixed number of cells that is highly noisy, increasing the number of clones in spots entangles the deconvolution problem. Consequently, for Tumoroscope-fixed, the highly noisy input confounds the model the most when the read counts are high and the model cannot assign the right clones to the spots, resulting in the largest MAE.

**Extended data Table 1:** The setups used for simulation.

| Property | Values |
| --- | --- |
| number of the clones in the evolutionary tree | 5 |
| number of mutations in the genotype | 30 |
| average clone per spot | 1, 2.5, 4.5 |
| average number of mutations per clone | 5.1, 13.6, 15 |
| average number of reads present in each spot | 18, 50, 80, 110 |
| average noise introduced to the number of cells | 0, 1, 10 |

**Extended data Figure 2:**
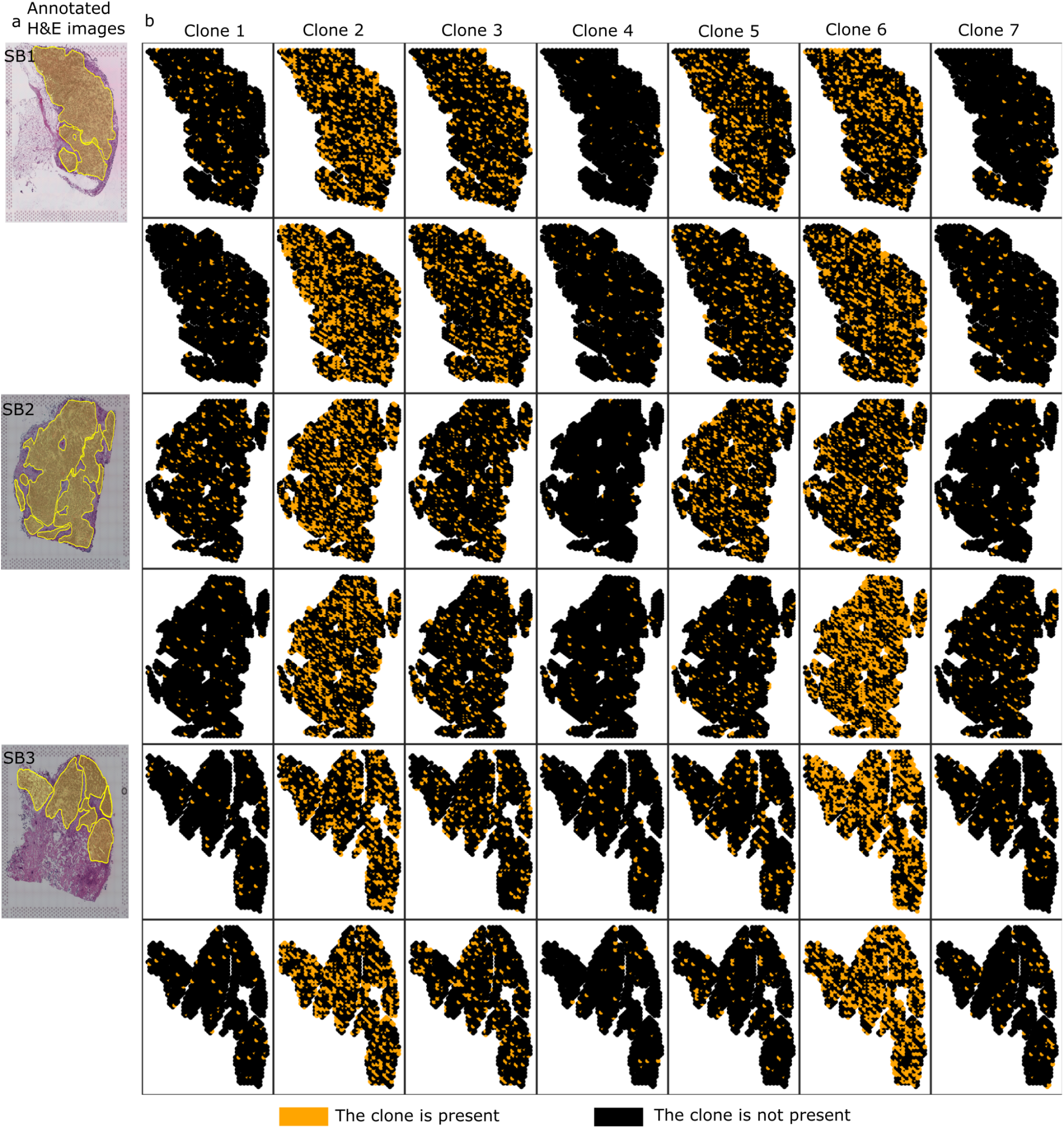
Spatial arrangement of cancer clones found by the cardelino for the breast cancer dataset. **a** Pathologist’s annotation of the cancerous areas on the H&E images for sections SB1, SB2, and SB3. **b** For each section, two rows correspond to the two nearby samples and seven columns correspond to the presence of the clone in the spots inferred by cardelino.

**Extended data Figure 3:**
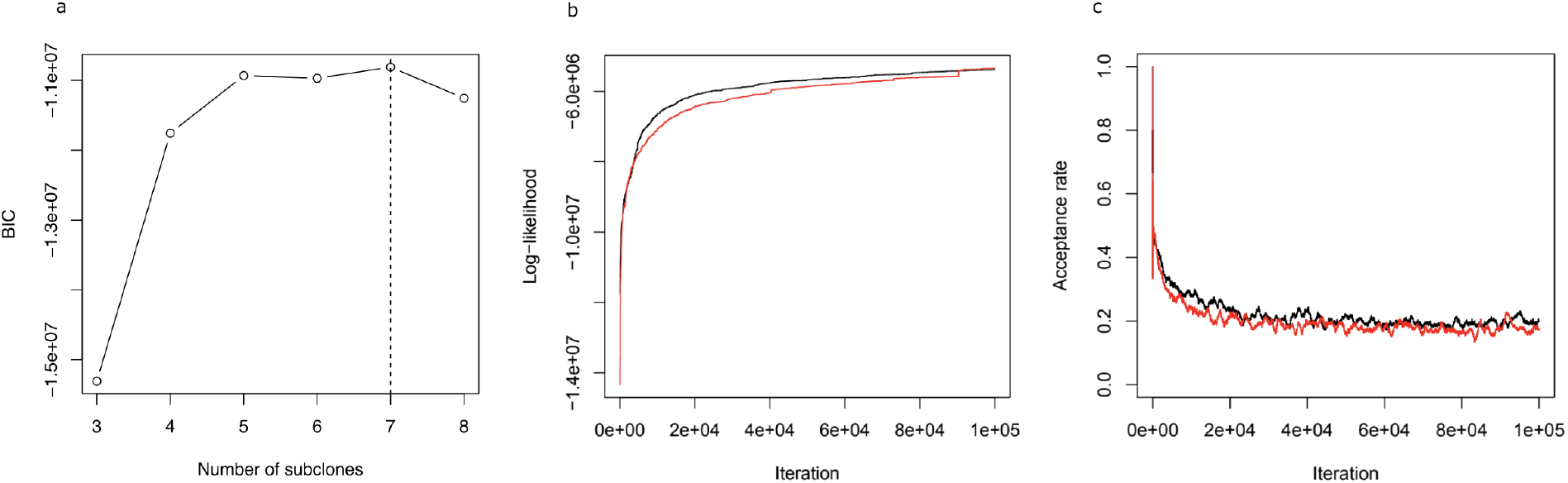
Analysis of the Canopy tree inference for the breast cancer dataset. **a** Bayesian Information Criterion (BIC; y-axis) of the Canopy model for different numbers of clones in the tree (x-axis). We selected the tree with seven clones, for which the BIC was the largest (indicated with the dotted vertical line). **b** Log-Likelihood of two MCMC chains of Canopy (y-axis) across MCMC iterations (x-axis), showing the convergence of the MCMC procedure. **c** Acceptance rate (y-axis) across iterations (x-axis). The acceptance rate converges to around the desired value of around 0.2.

**Extended data Figure 4:**
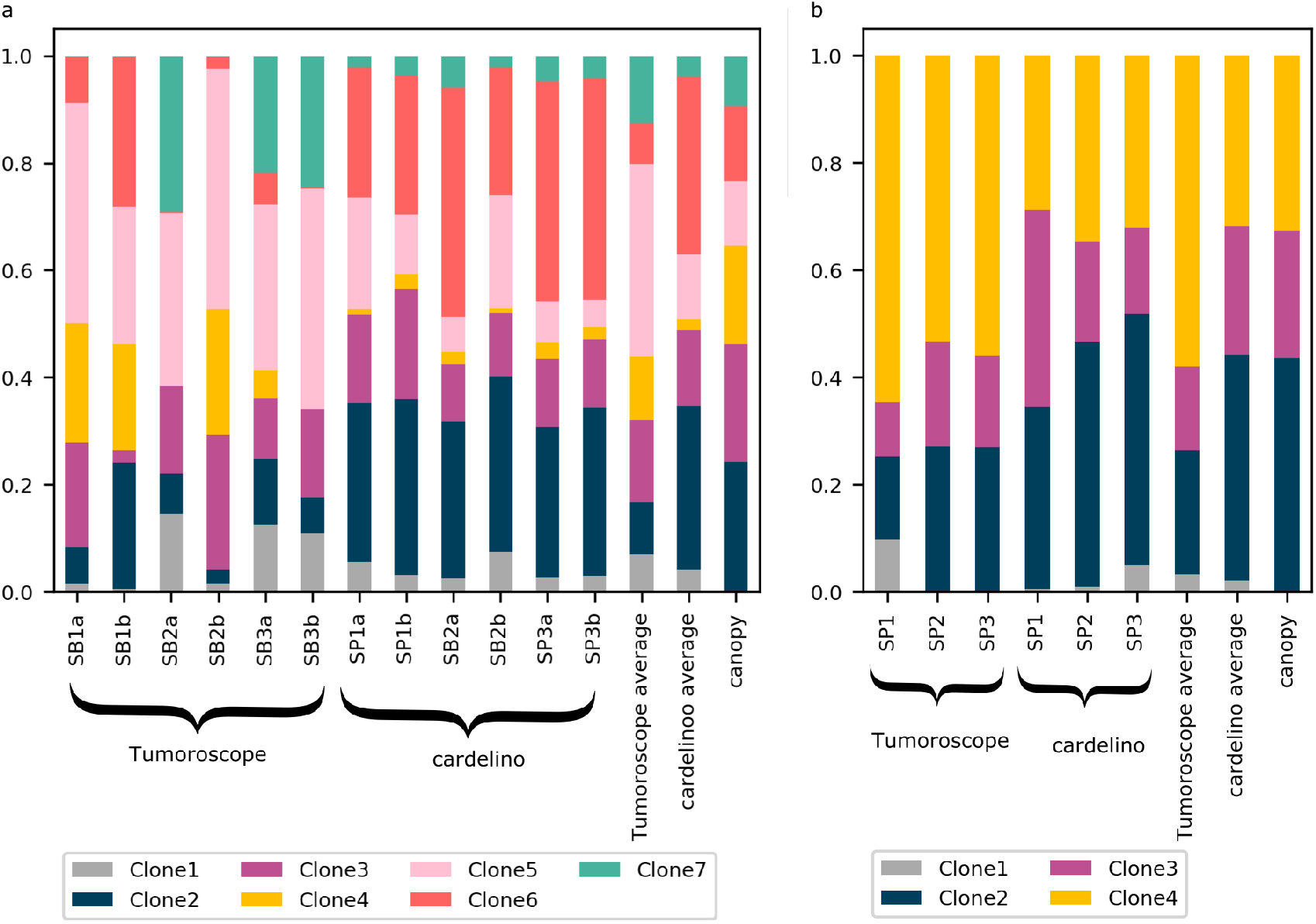
Proportion of each clone in each section. **a** Proportions of inferred clones by Tumoroscope and cardelino for the breast cancer dataset. The proportions were computed by summing the inferred fractions across spots for each ST section. Averages over sections and clone frequences inferred by Canopy from bulk DNAseq data are also shown. **b** Proportions of inferred clones by Tumoroscope and cardelino for the prostate cancer dataset.

**Extended data Figure 5:**
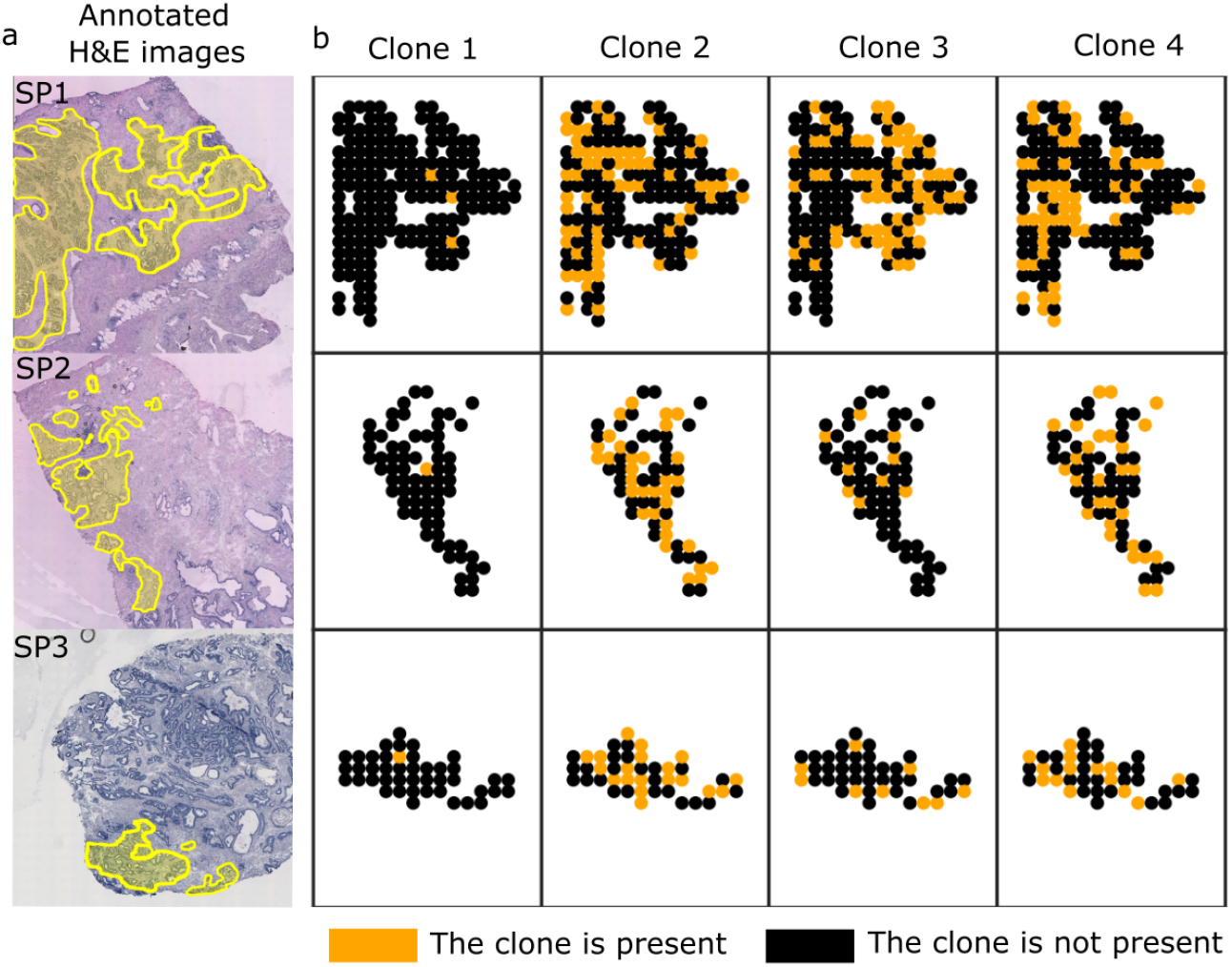
Results obtained by cardelino for the prostate cancer dataset. **a** Pathologist’s annotation of the cancerous areas on the H&E images for sections SP1, SP2, and SP3. **b** For each section (rows), there are four columns corresponding to the presence of the clones in the spots.

**Extended data Figure 6:**
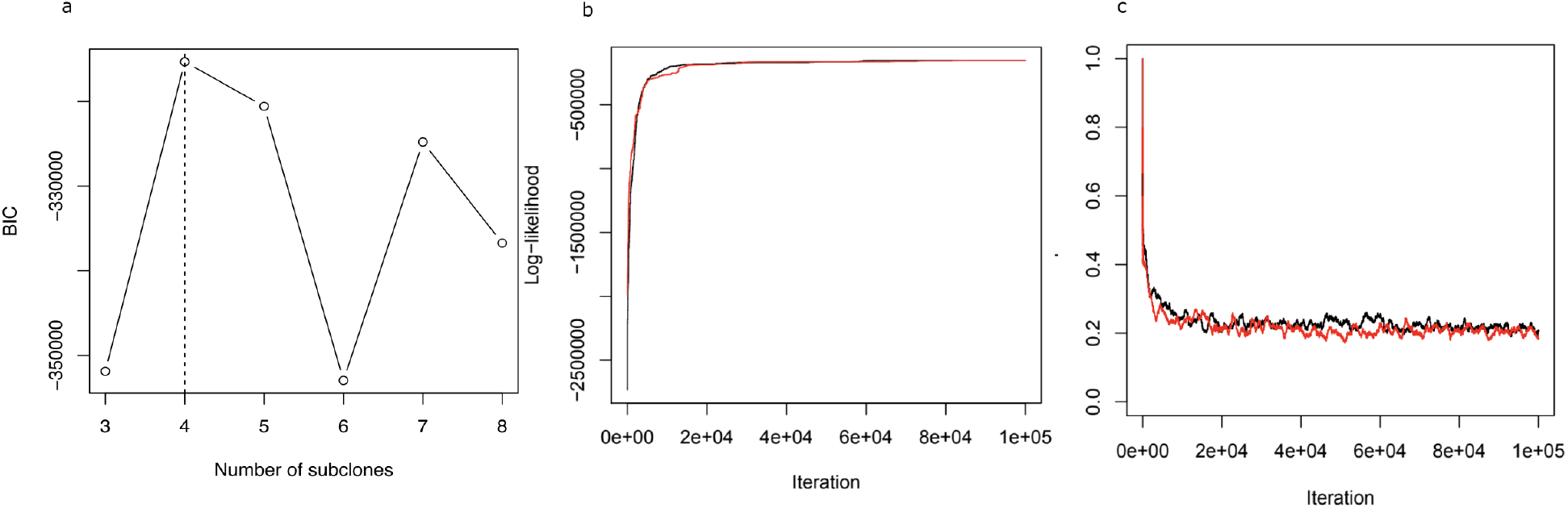
Analysis of the Canopy tree inference for prostate cancer dataset. **a** Bayesian Information Criterion (BIC; y-axis) of the Canopy model for different numbers of clones in the tree (x-axis). We selected the tree with four clones, for which the BIC was the largest (indicated with the dotted vertical line). **b** Log-Likelihood of two MCMC chains of Canopy (y-axis) across MCMC iterations (x-axis), showing the convergence of the MCMC procedure. **c** Acceptance rate (y-axis) across iterations (x-axis). The acceptance rate converges to around the desired value of around 0.2.

**Extended data Figure 7:**
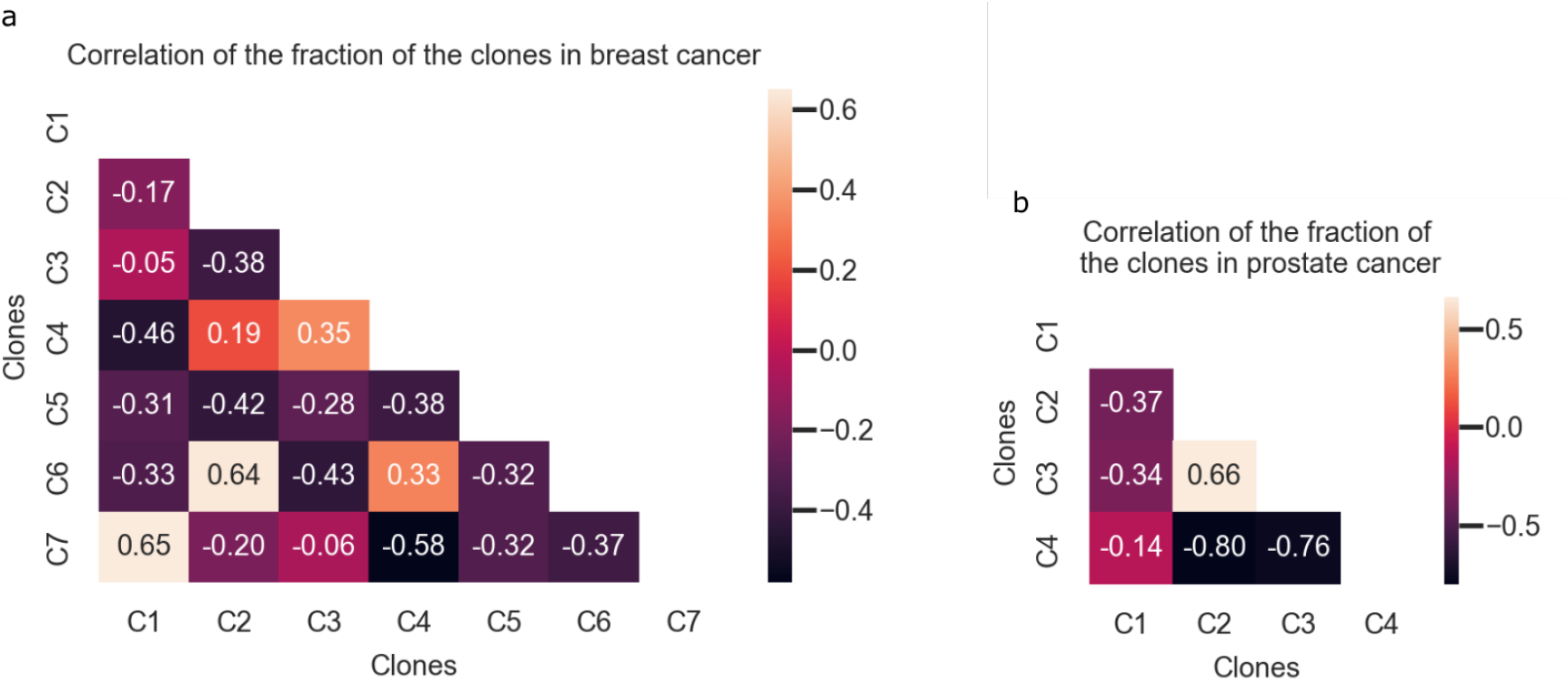
Pairwise Pearson correlations of the proportions of all the spots taken by the clones for breast cancer (**a**) and for prostate cancer (**b**) data.

**Extended data Figure 8:**
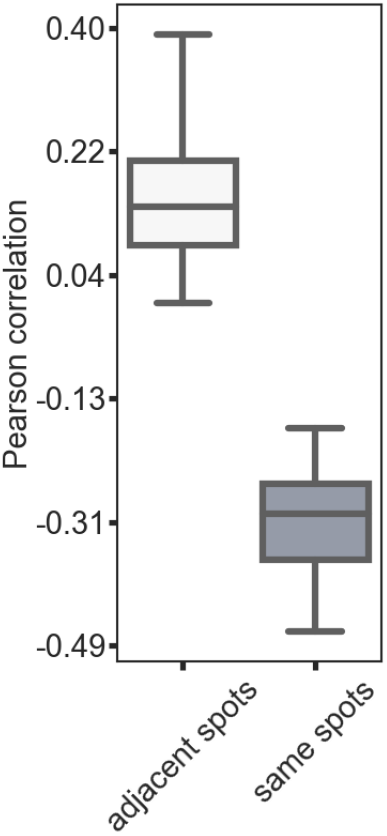
Distribution of the Pearson correlation between the proportions of the spots taken by the clones 3 and 5. The correlation between proportions of clone 5 and 3 in adjacent spots was computed for 20 different sets of randomly sampled pairs of 100 adjacent spots. The correlation between proportions of clone 5 and 3 in the same spots was computed for 20 sets of 100 randomly sampled spots.

